# The Connectome Modulates Critical Brain Dynamics Across Local and Global Scales

**DOI:** 10.64898/2025.12.11.693658

**Authors:** Giovanni Rabuffo, Pietro Bozzo, Bach Nguyen, Damien Depannemaecker, Marco N. Pompili, Leonardo L. Gollo, Tomoki Fukai, Pierpaolo Sorrentino, Leonardo Dalla Porta

## Abstract

Neuronal activity in the brain has been hypothesized to operate near criticality–a dynamical regime poised between order and disorder that maximizes information processing, adaptability, and dynamic range. While criticality has been extensively studied at local scales (within neuronal populations) and at global scales (across interacting brain regions), the interplay between these levels remains poorly understood. Here, we propose a multiscale computational framework that bridges *local* and *global* criticality within a single, mechanistic model. At the mesoscopic level, individual brain regions are represented by neural mass models tuned near the transition between asynchronous and synchronous regimes. These regions are then coupled via an empirically derived mouse connectome to investigate how structural connectivity shapes the emergence of large-scale coordination. We show that (i) local near critical dynamics for an isolated brain region can be faithfully reproduced within a mean-field model framework, (ii) local distance to criticality is modulated by long-range coupling, (iii) whole-brain simulations reveal non-linear gradients of timescales and heterogeneous shifts towards/away from local criticality, and (iv) global criticality, manifested in scale-free avalanche distributions and optimal functional connectivity, emerges when local populations are locally tuned near criticality and coupled within an optimal range. These results demonstrate that local and global criticality are dynamically intertwined but not directly aligned, and that their relationship depends on the underlying structural connectivity. Our multiscale modeling framework provides a tractable tool for generating testable hypotheses on how brain criticality co-arises across scales and how it may be modulated in health and disease.

## I Introduction

The concept of brain criticality–that the brain operates near a phase transition between order and disorder–has become a central framework in theoretical and empirical neuroscience [1–5]. At a critical point, neuronal activity exhibits long-range correlations [6], scale-free avalanches [7], and optimal computational properties such as maximal dynamic range and information transmission [8–10]. Evidence for critical-like behavior has been found both in local regions [11, 12], where it emerges from the interactions among neurons within a cortical patch or microcircuit, and at the global scale [13–20], where it reflects coordination between distant brain regions (for a recent review see [5]). Yet, these two levels of description have largely been studied separately.

From a theoretical standpoint, scale invariance suggests that the same principles could govern dynamics across spatial scales [21]. However, many brain features are specific to a particular scale, as opposed to being scale-free [22, 23]: distinct structural features characterize different levels, from dense, recurrent microcircuits to sparse, heterogeneous long-range connections. As a result, criticality may not simply ‘propagate’ across scales. Local circuits tuned near criticality might not imply criticality at the whole-brain level, and conversely, large-scale coordination may emerge even when local dynamics deviate from criticality. Understanding this multiscale relationship is key to linking cellular and systems-level theories of brain function.

Here, we develop a multiscale modeling formalism to study local and global criticality at once (Fig. 1A). At the mesoscopic level, each brain region is described by an all-to-all coupled network of ‘active rotator’ neurons [24], which reproduces subcritical, critical, and supercritical regimes [25]. This network admits an analytic mean-field reduction [25, 26], allowing us to systematically characterize regional dynamics and their dependence on local parameters. At the macroscopic level, distinct brain regions are coupled according to a mouse structural connectome [27, 28], enabling us to study how anatomical connectivity shapes large-scale coordination and modulates each region’s proximity to criticality.

**FIG. 1.**
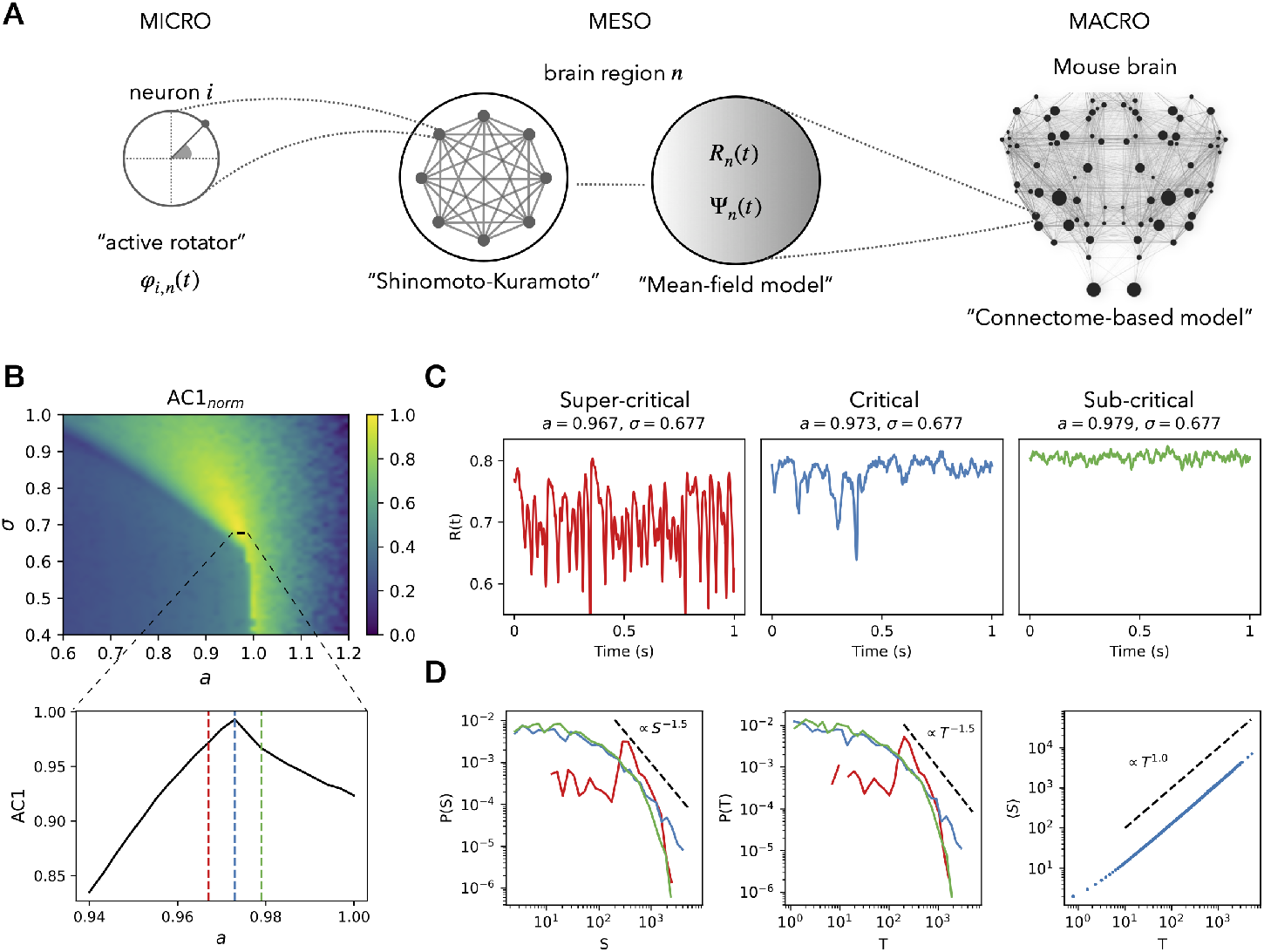
Multiscale modeling and criticality analysis of brain region dynamics. **A)** Schematic of the multiscale framework. At the *micro* scale, each neuron *i* in the brain region *n* is modeled as an active rotator *φ*_*i,n*_(*t*). At the *meso* scale, the collective dynamics are described by the Shinomoto–Kuramoto model for a population of all-to-all connected active rotators. This model admits a mean-field description, which reduces the neuronal population dynamics to only two equations for the mesoscopic variables (*R*_*n*_(*t*), Ψ_*n*_(*t*)). When driven by uniformly distributed noise of amplitude *N* = 0.0001, the mean-field model describes the activity of an isolated brain region. At the *macro* scale, brain regions are coupled according to a mouse connectome from the Allen Institute [27] **B)** Exploration of the local model’s parameter space. The heatmap shows the normalized autocorrelation at lag 1 ms (AC1) of *R*(*t*) over different values of the coupling parameter *a* and noise amplitude *σ*, revealing a ridge of critical-like dynamics. A zoom-in along *σ* = 0.677 highlights representative supercritical (red; *a* = 0.967), critical (blue; *a* = 0.973), and subcritical (green; *a* = 0.979) regimes, consistent with microscopic network simulations [25]. **C)** Example time series of *R*(*t*) in each near-critical regime. **D)** Avalanche analysis: avalanches are defined as consecutive time points where *R*(*t*) exceeds its median value. Shown are distributions of avalanche sizes *S* (defined as the area between the curve and the threshold) and durations *T*, as well as the scaling of average size ⟨*S* ⟩ versus duration *T*. Straight dashed lines mark theoretical power laws consistent with critical dynamics in the blue (critical) regime.

We first show that–using only two coupled, noise-driven variables–the local neural mass model reproduces hallmark signatures of criticality, including increased temporal correlations and scale-free avalanche statistics. We then investigate how long-range coupling between regions in different regimes can shift the dynamics of the downstream region toward or away from criticality, revealing non-trivial cross-regime influences. Extending to whole-brain simulations, we show that global coupling systematically modulates local regimes according to the region-specific connectivity, giving rise to topological gradients of timescales whose direction depends on whether regions are tuned to subcritical, critical, or supercritical states. Crucially, we find that global criticality–reflected in large-scale avalanche scaling and optimal fit to empirical functional connectivity–emerges when all local populations are tuned near criticality and weakly coupled.

These results advance a unifying view in which local and global criticality are not independent phenomena but dynamically coupled aspects of brain organization. By embedding minimal models of local dynamics into a realistic connectome, our approach bridges levels of description and provides a tractable testbed for investigating how structural connectivity and dynamical tuning interact to shape multiscale coordination. Beyond its theoretical significance, this framework offers concrete hypotheses for experimental probing of criticality across scales, and may help explain how pathological alterations in local or global dynamics disrupt brain function.

## II Results

### A A mean-field description of local critical dynamics

We first investigated whether a reduced mean-field model formulation could reproduce the critical-like dynamics observed in detailed neuronal network models. We modeled each isolated brain region as the mean-field limit of an all-to-all connected population of active rotators [25, 26], retaining the key Kuramoto mesoscopic variables (*R*_*n*_(*t*), Ψ_*n*_(*t*)), representing the population’s synchronization and average phase, while achieving a substantial reduction in dimensionality (Fig. 1A). Adding stochastic noise and exploring the model parameters, including the network excitability *a* and the noise amplitude *σ*) (see methods for details), we generated non-trivial meanfield dynamics for a population of neurons (Suppl. Fig. S1 A-B).

Near a critical point, neural populations exhibit critical slowing down, meaning that perturbations decay more slowly and activity becomes increasingly temporally correlated [29]. This makes short-lag autocorrelation a sensitive indicator of proximity to criticality [30]. Accordingly, we quantified temporal structure using the autocorrelation function at a 1-ms delay (AC1). Systematic exploration of the parameter space revealed a ridge of critical-like dynamics, where AC1 is maximized (Fig. 1B-C, blue).

Moving from this critical regime, fixing *σ* = 0.677 and decreasing the parameter *a* produces oscillatory dynamics, corresponding to a supercritical regime with highly synchronous neuronal activations (Fig. 1B-C, red). Conversely, increasing *a* leads to a subcritical regime with asynchronous, low-activity dynamics (Fig. 1B-C, green). Note that in the latter case, the Kuramoto order parameter *R*(*t*) remains high, as a result of prolonged silent periods, reflecting synchrony in weak neuronal activity (Suppl. Fig. S2). Neural network simulations by Buendia and colleagues show a similar phase diagram for a network of all-to-all connected active rotators (see the hybrid-type regimes in Figure 3 of [25]). The authors demonstrate diverse bifurcation points between synchronous and asynchronous states, with the critical regime corresponding to a hybrid-type (HT) transition, where incipient oscillations coexist with an asynchronous state. Running deterministic simulations, we confirmed the existence of bistability in the mean-field model for the parameters that maximize the AC1 (Suppl. Fig. S1 C).

Avalanche analyses confirmed that, at the HT transition, the mean-field model reproduces scale-free avalanche size and duration statistics consistent with critical dynamics (Fig. 1D). These results validate the minimal model as a useful representation of near-critical local neuronal population dynamics, suitable for large-scale coupling in connectome-based simulations.

### B Cross-regime coupling can shift local dynamics toward or away from criticality

To examine the mechanistic basis of how long-range coupling influences local regimes, we simulated unidirectionally connected pairs of brain regions *A* (source) and *B* (target) exploring the (*a*_*A*_, *a*_*B*_) parameters systematically. Depending on the initial dynamical states, the activity in the target region varied. For example, when a supercritical source was coupled to a subcritical target (Fig. 2A-B), or vice versa (Fig. 2C-D), the target region shifted toward the critical regime. Coupling two critical brain regions did not alter the dynamical state of the target, preserving criticality (Fig. 2E-F). Conversely, coupling a subcritical source to a critical or subcritical target further increased the target’s distance from criticality (not shown). These results are summarized in Fig. 2G, reporting the normalized difference ΔAC1 between AC1 of population B in coupled versus uncoupled cases. Increased or decreased AC1 marks an approach to or a distancing from criticality, respectively. This interpretation follows directly from the single-population results (Fig. 1B), where AC1 exhibited a pronounced peak along the critical ridge. There, the maximized short-lag autocorrelation reflected critical slowing down, providing a sensitive indicator of proximity to the transition. By analogy, changes in AC1 in the coupled-population model can therefore be taken as shifts toward or away from this critical regime. When the populations are mutually coupled with feedback loops (Fig. 2H), results are comparable to the only feed-forward case, and differ mostly when the upstream population dwells around a critical state (Fig. 2I), demonstrating higher sensitivity in these regimes. In summary, the long-range coupling can alter the local dynamical regime of a brain region, promoting or disfavoring critical dynamics.

**FIG. 2.**
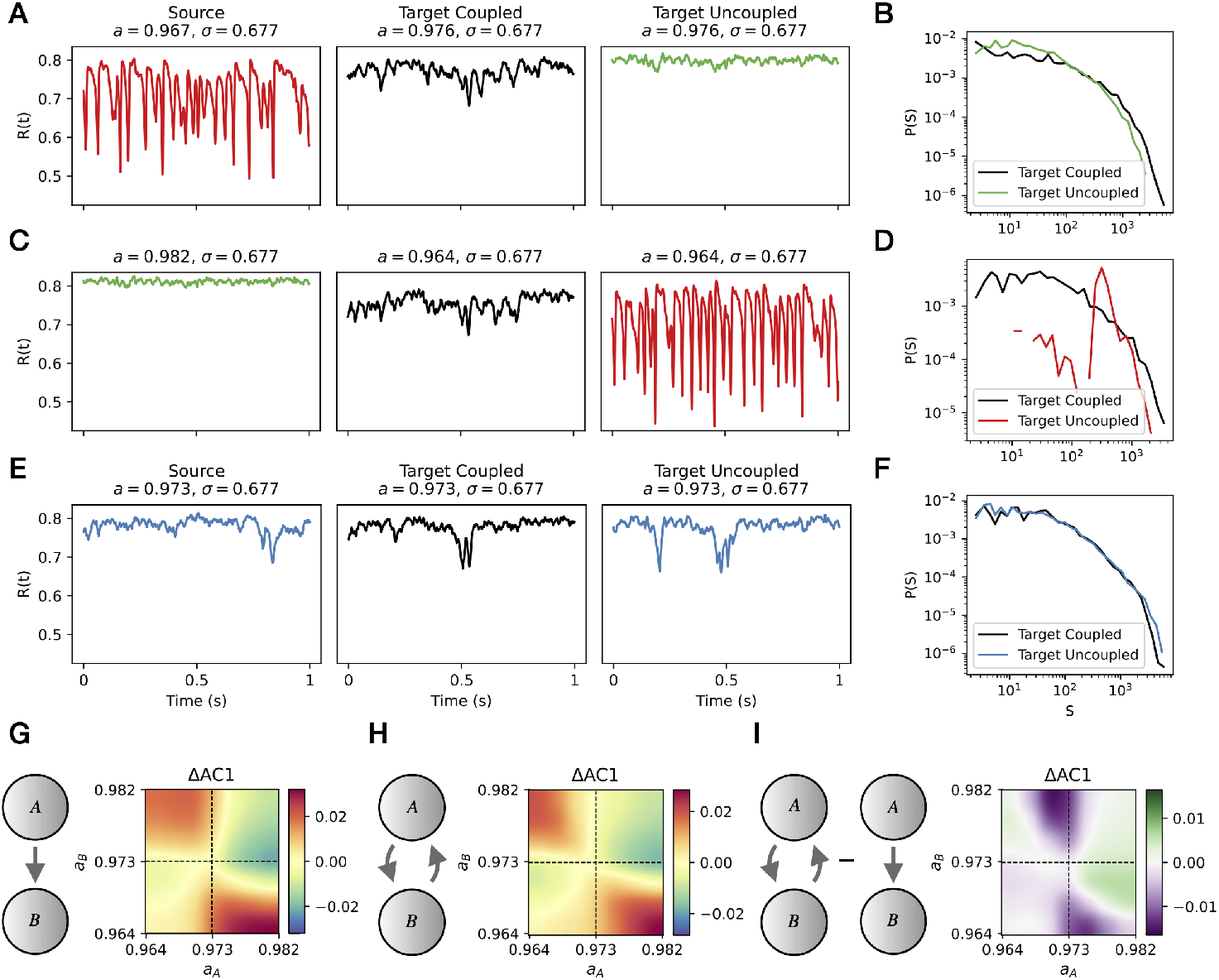
Coupling between brain regions modulates criticality in the target. **A–F)** Two mesoscopic populations, A (source) and B (target), are coupled unidirectionally. We fix the parameter *σ* = 0.677 as in Figure 1, and the global coupling to *G* = 0.05 (for details on the coupling function refer to the Methods section). We systematically vary the parameters (*a*_*A*_, *a*_*B*_) of each region to place them in supercritical, critical, or subcritical regimes, and assess the effect of coupling on the target population B. Example time series and avalanche distributions for three representative cases: **A–B)** supercritical source to subcritical target, **C–D)** subcritical source to supercritical target, and **E–F)** critical-to-critical coupling. In each case, coupling alters the target dynamics and modifies its avalanche statistics, often enhancing scale-free properties near criticality. **G)** Heatmaps show the relative change of AC1 computed on the target’s dynamics (defined as ΔAC1 = (AC1_B,coupled_− AC1_B,uncoupled_)*/*AC1_B,uncoupled_). The results demonstrate that cross-regime coupling can induce non-trivial effects, often driving the target closer to criticality when source and target begin in contrasting dynamical states (e.g., supercritical → subcritical, or vice versa). **H)** Similar to panel G, for mutually coupled populations. **I)** Relative difference between panels H and G.

### C Critical local tuning best matches empirical FC and dFC at intermediate coupling

We simulated whole-brain dynamics by coupling neural mass models through the directed structural connectivity of the Allen Mouse Brain Atlas [27, 28]. Each brain region is represented as a network node, and nodes are initialized homogeneously in one of three dynamical regimes–subcritical, critical, or supercritical–depending on the choice of local parameters (i.e., keeping *σ* = 0.677 fixed, and using the three values of parameter *a* as in Fig. 1). The coupling strength between regions is controlled by a global parameter *G*, which scales the influence of long-range connectivity on local dynamics: for *G* = 0, regions evolve independently, while increasing *G* progressively introduces inter-regional interactions. In the next sections, we investigate the results of whole-brain simulations at two levels: 1) using the fast neuroelectric signals at 2000 Hz (i.e., the same type of signal analyzed in the previous sections); and 2) using the slow BOLD signals at 1 Hz, obtained by applying a Balloon–Windkessel hemodynamic response function (HRF) to the neuroelectric signals.

First, to ground our simulations to empirical data, we compared the simulated BOLD activity with resting-state fMRI recordings from 53 control mice under light anesthesia (Fig. 3A). For each mouse recording, we evaluated model performance by comparing simulated and empirical functional connectivity (FC) and dynamic FC (dFC) statistics across a range of *G* (Fig. 3B). For all three local regimes, an appropriate choice of *G* reproduced empirical FC and dFC distributions. The optimal working point *G*^∗^ depended systematically on the local regime: the *critical* model matched the data at low coupling (*G*^∗^ ≈ 0.03), the *supercritical* model required a intermediate coupling (*G*^∗^≈ 0.07), and the *subcritical* model achieved comparable fits only at higher coupling values (*G*^∗^ ≈ 0.21). This ordering is consistent with the higher sensitivity of critical neural masses, for which weaker interregional coupling suffices to coordinate large-scale dynamics [31]. Example simulated BOLD rasters, FC matrices, and dFC matrices at each regime’s working point (Fig. 3C) illustrate the corresponding qualitative differences in functional structure and variability.

**FIG. 3.**
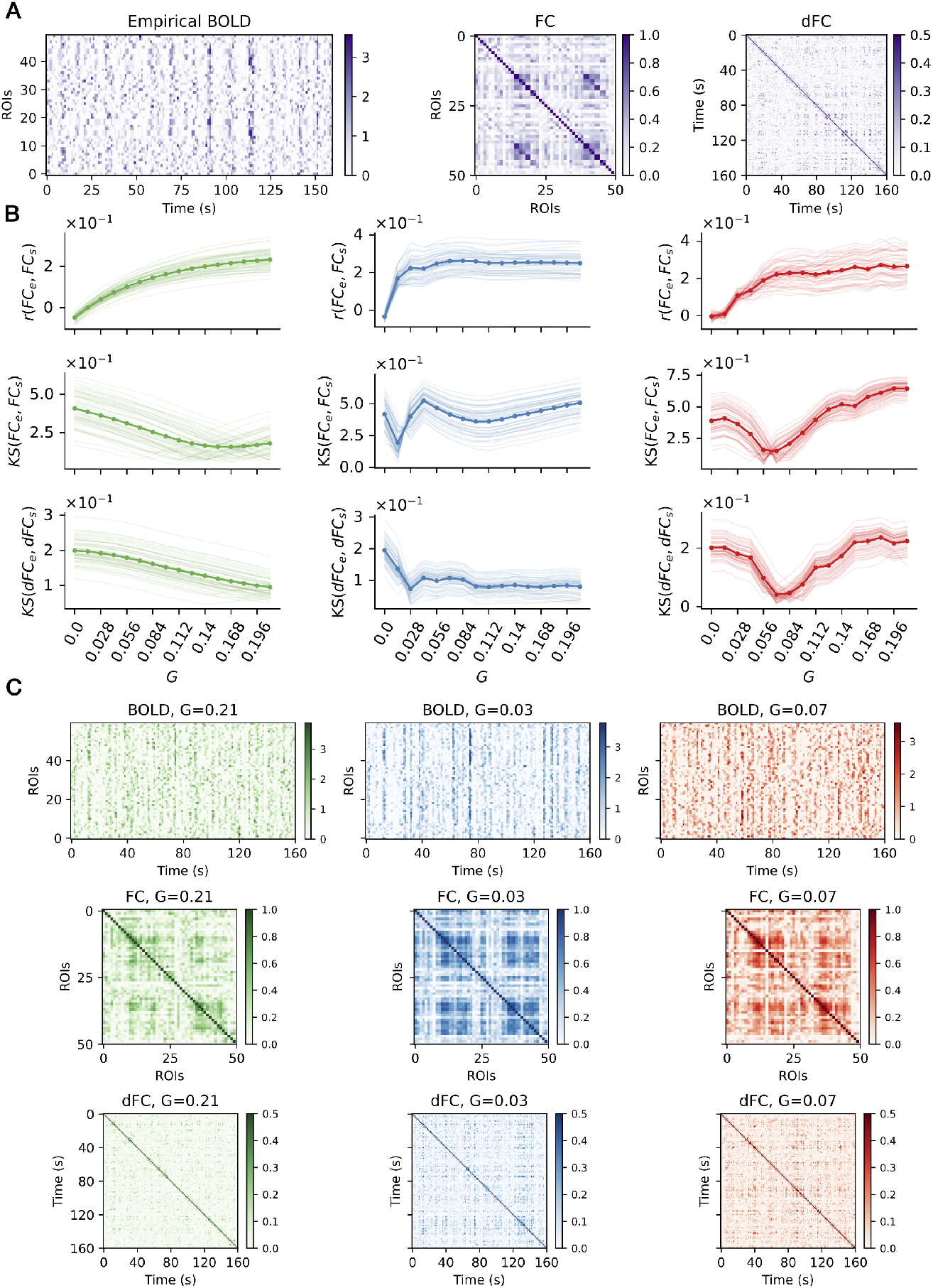
Model performance against empirical fMRI data across dynamical regimes. **A)** Empirical fMRI BOLD signal (left), functional connectivity (FC; middle), and dynamic FC (dFC; right) from an example mouse. **B)** Comparison between simulated and empirical FC and dFC statistics across a range of global coupling values *G*, for subcritical (green), critical (blue), and supercritical (red) regimes. For each of these regimes, local parameters *a* and *σ* are fixed homogeneously across all ROIs as in Figure 1C. Each thin line represents one mouse. The thick line represents the average across mice. Top: correlation between simulated and empirical FC as a function of *G*. Middle: Kolmogorov–Smirnov (KS) distance between simulated and empirical FC distributions. Bottom: KS distance between simulated and empirical dFC distributions. In all metrics, the critical regime (blue) yields the best match to empirical data at intermediate *G* values. **C)** Example simulated BOLD rasters (top), FC matrices (middle), and dFC matrices (bottom) at the optimal coupling *G* for each regime, illustrating qualitative differences in functional structure and variability.

### D Global criticality emerges at best fit of empirical data

We quantified how local tuning and long-range coupling shape *global* network signatures of criticality at the level of fast neuroelectric dynamics while initializing each region *n* in a common local regime (subcritical, critical, or supercritical; Fig. 4A). At the optimal working point (i.e., the best fit to BOLD empirical data, Fig. 3C), all three configurations displayed the emergence of coordinated large-scale bursts, which can be visualized as quasi-instantaneous high-amplitude cofluctuations in the raster plots in Figure 4B.

**FIG. 4.**
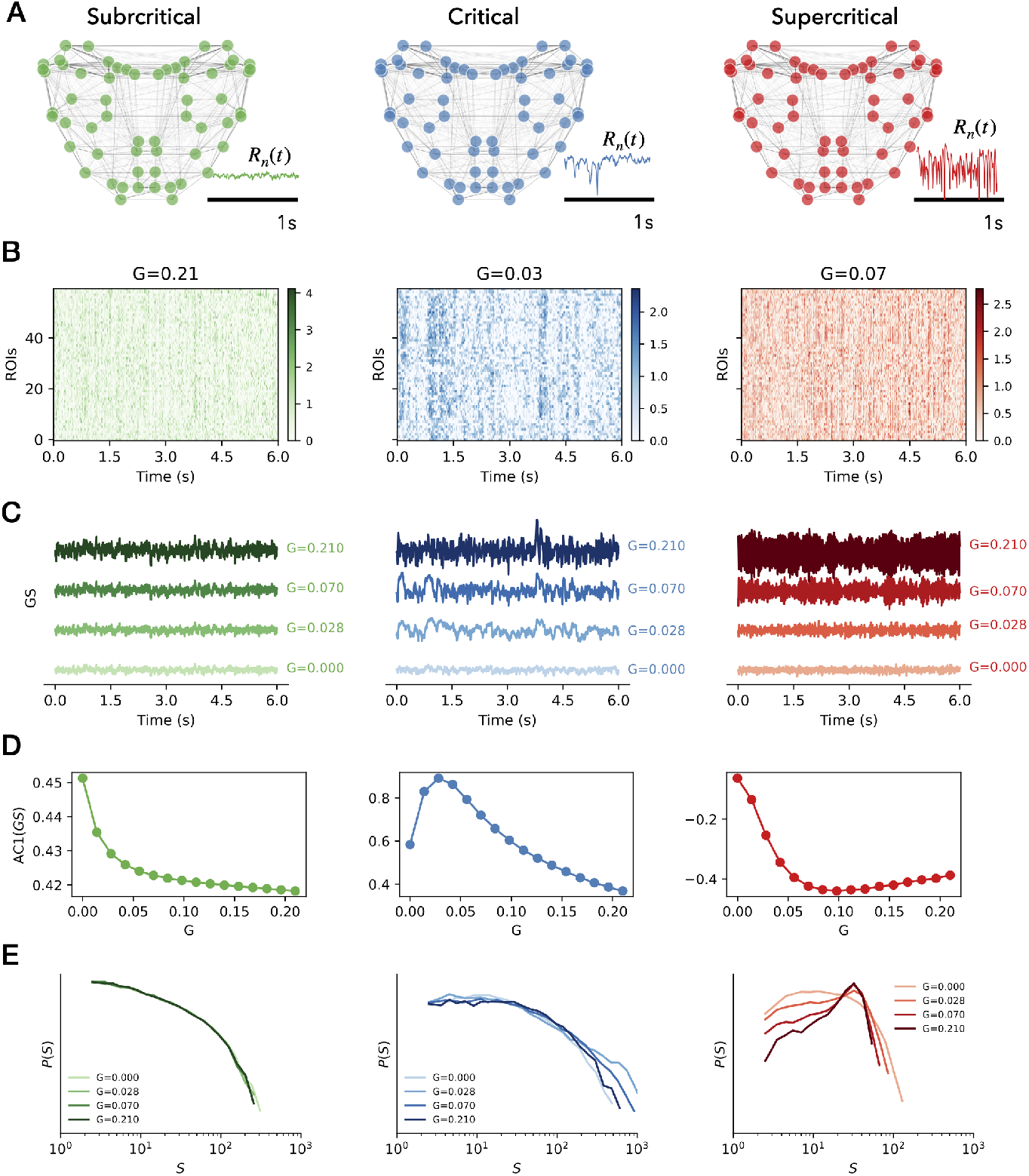
Global signatures of criticality across whole-brain simulations. **A)** Whole-cortex simulations of fast neuroelectric timeseries using the mouse structural connectome. All brain regions are initialized in either a subcritical (green), critical (blue), or supercritical (red) regime, and coupled via the anatomical connectivity. Example time series *R*_*n*_(*t*) illustrate the resulting local dynamics for each regime. **B)** Raster plots showing standardized neuroelectric activity across all brain regions (ROIs) for three working point values of the global coupling strength *G*, under subcritical (green), critical (blue), and supercritical (red) local tuning. **C)** Corresponding global signals, computed as the average across all standardized ROI activities. As *G* increases, the amplitude and temporal structure of the global signal change in regime-specific ways. **D)** Lag-1 autocorrelation (AC1) of the global signal *GS*(*t*) as a function of *G*. In the subcritical regime (green), AC1 decreases monotonically with *G*, indicating increasing desynchronization. In the critical regime (blue), AC1 peaks at intermediate *G*, revealing an optimal coupling range where global dynamics are closest to criticality. This optimal range coincides with the working point where simulations match the empirical data (Fig.3). In the supercritical regime (red), AC1 increases and then saturates, consistent with dominance of strong local oscillations. **E)** Distributions of global avalanche sizes *S*, defined from threshold crossings of the global signal. The emergence of power-law-like scaling occurs most prominently in the critical case (blue) at intermediate *G*, indicating the onset of scale-free, globally coordinated dynamics.

Starting from *G* = 0 and gradually increasing the global coupling, we inspected the effects of long-range connectivity on the corresponding global signal *GS*(*t*), defined as the average across all standardized ROI activities, 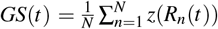, (Fig. 4C). Although stronger coupling promotes spatial coordination across regions in all three regimes– reflected in increased *GS* amplitude at higher *G* values–the temporal structure of *GS*(*t*) changed in regime-specific ways (Fig. 4C): subcritical tuning produced progressively faster fluctuations; critical tuning amplified slow, aperiodic fluctuations over an intermediate *G* range; and supercritical tuning led to large-amplitude, quasi-oscillatory fast bursts at higher *G*.

To quantify these changes, we measured the AC1 of the global signal as a function of *G* (Fig. 4D). In the *subcritical* regime (green), AC1 decreased monotonically with *G*, indicating that stronger recurrent coupling accelerated the relaxation of the population-average signal and thereby reduced its temporal persistence. In the *critical* regime (blue), AC1 displayed a pronounced, non-monotonic dependence on *G*, peaking at intermediate coupling–an operating range in which inter-regional interactions optimally prolong global timescales. In the *supercritical* regime (red), AC1 decreased and displayed a minimum at intermediate *G* values, reflecting a nontrivial interaction between local oscillations and variability in their synchronization.

We then probed global coordination through an avalanche analysis of *GS*(*t*) (Fig. 4E). Global avalanches were defined as excursions of *GS*(*t*) above a threshold (the median of the timeseries). Avalanche sizes *S* were defined as the total excursion area during each event. Power-law-like scaling in the distribution *P*(*S*) emerged most prominently for *critical* local tuning at intermediate *G*, signaling the onset of scale-free, system-wide coordination. In contrast, subcritical networks produced steeper, truncated distributions (reflecting weak, rapidly damped excursions), whereas supercritical networks developed heavy tails at large *S* but with deviations from scale-free structure due to strongly driven, large-amplitude events.

Together, these findings indicate that *global* criticality is not achieved by coupling alone, but rather emerges from the interaction between local near-critical dynamics and intermediate coupling. Remarkably, signatures of global criticality obtained using the fast neuroelectric signals–such as the maximization of *AC*1(*GS*) (Fig. 4D) and of the power-law-like scaling range of global avalanches (Fig. 4E)–emerged when local neural masses were tuned to criticality and the global coupling was set to the working point (*G* = 0.028) that best reproduced empirical mouse fMRI data (thus, at the BOLD level; Fig. 3B–C). This alignment links global critical signatures to improved model–data correspondence.

### E Long-range interactions produce heterogeneous shifts in criticality

Although regions are homogeneously initialized in a common local regime (subcritical, critical, or supercritical), long-range coupling drives the neural masses away from their initial state, consistent with our analysis in Fig.2. We quantified how varying the coupling strength *G* modulates fast regional dynamics by measuring the AC1 of each simulated neuroelectric signal *R*_*n*_(*t*) at coupling *G* (Fig. 5A). For each region *n*, we related the timescale AC1(*R*_*n*_(*t*)) emerging from large-scale coupling to its structural in-strength (i.e., the sum of incoming structural weights; Fig. 5B).

**FIG. 5.**
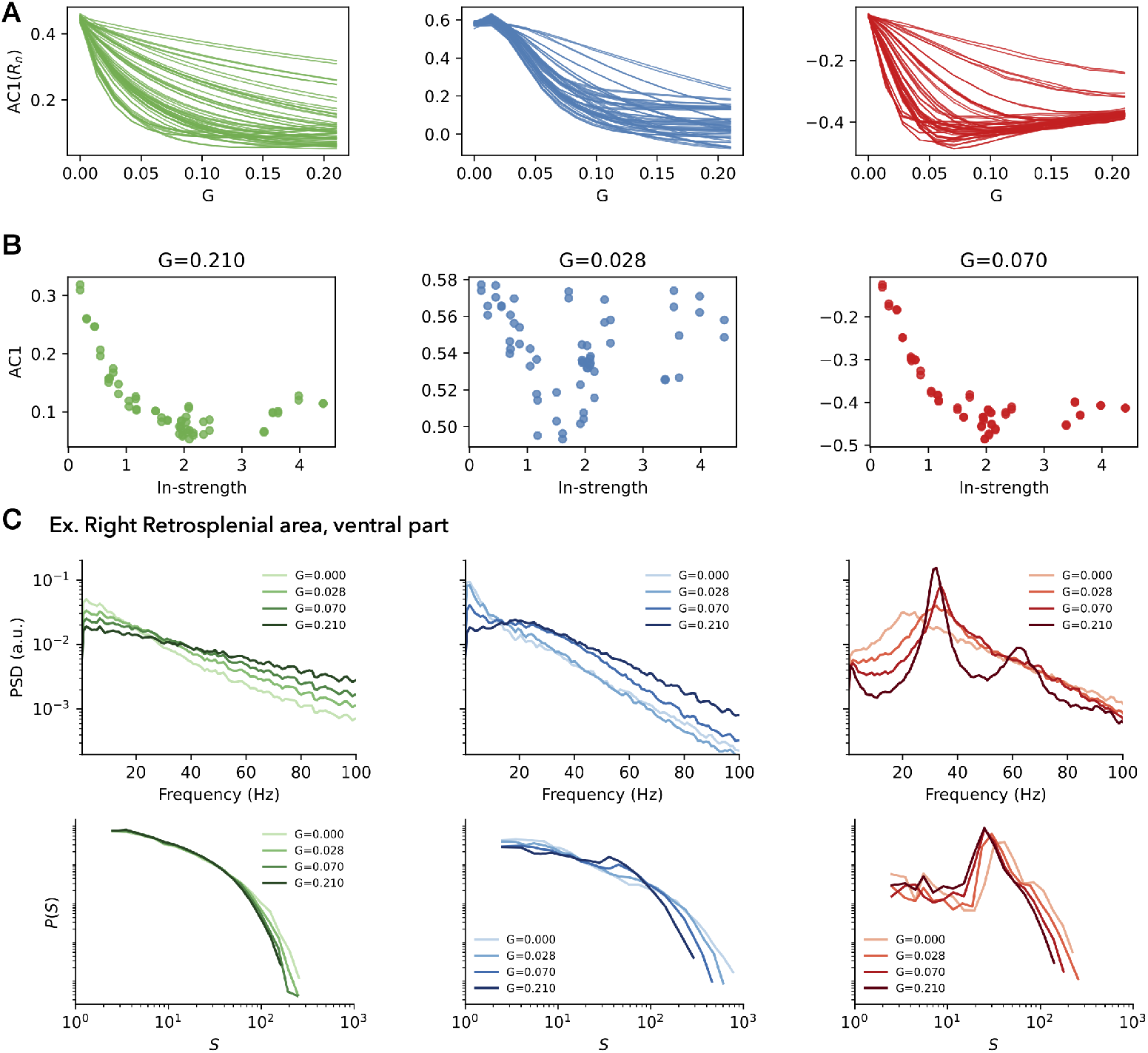
Connectome-based simulations reveal interaction between local criticality and global coupling. **A)** Effect of increasing the global coupling strength *G* on the lag-1 autocorrelation (AC1) of regional neuroelectric activity (one line per region). In the subcritical case (green), coupling monotonically decreases AC1, driving regions further from criticality. In the critical case (blue), AC1 initially increases with *G*, pushing regions closer to criticality before decreasing again. In the supercritical case (red), the effect of coupling is non-monotonic and heterogeneous across regions. **B)** Change in AC1 relative to the uncoupled case (ΔAC1) plotted against each region’s in-strength (sum of incoming structural connectivity weights). In the subcritical and critical regimes, AC1 decreases with structural in-strength, reflecting a smooth gradient. In the critical regime, this relation is lost and non-linear effects are more marked. **C)** Example dynamics from the right retrosplenial area (ventral part). Power spectral density (PSD) shows characteristic spectral changes across regimes: 1*/ f* steepening in the subcritical case, suppression of slow frequencies in the critical case, and emergence of fast oscillations in the supercritical case. Avalanche size distributions for the same region further reveal that in the critical condition (blue), increased coupling slightly shifts dynamics toward a supercritical regime at higher *G*.

Across different starting regimes, coupling induces distinct, topology-dependent patterns: (i) *Subcritical regime (green):* for most regions, AC1 decreases monotonically with *G*, shifting dynamics away from criticality (Fig. 5A). This decrease exhibits a pronounced topological gradient: nodes with higher in-strength showed smaller AC1, consistent with faster timescales and stronger afferent drive accelerating local relaxation (Fig. 5B). (ii) *Critical regime (blue):* the highest AC1 is observed for weak *G*, bringing regions closer to criticality, but decreases at higher coupling *G* (Fig. 5A). The relationship between AC1 and in-strength is strongly non-linear: nodes with intermediate in-strength exhibit faster timescales, whereas under strong coupling the system reverts to slower timescales (Fig. 5B). (iii) *Supercritical regime (red):* across coupling values, AC1 is predominantly negative, reflecting oscillatory dynamics rather than slow fluctuations (Fig. 5A). Here, the relationship with in-strength is again monotonic; however, compared to the subcritical case, the timescales increase in magnitude, indicating stronger oscillatory persistence in regions with higher in-strength. (Fig. 5B).

These regime-specific signatures are mirrored in the spectral statistics of exemplar regions (Fig. 5C, top). In the right retrosplenial area–a functional hub of the mouse connectome [32]–the power spectral density reveals distinct spectral signatures across regimes. In the subcritical state, increasing global coupling flattens the 1*/ f* slope. At criticality, when global coupling is tuned to the working point that best matches empirical data and yields global criticality (*G* = 0.028, Fig. 3 -4) local activity exhibits a canonical 1*/ f* decay. In the supercritical regime, coupling introduces additional high-frequency components, reflecting the emergence of new rhythms through long-range interactions. Consistently, local avalanche size distributions in the retrosplenial area (Fig. 5C, bottom) indicate that local criticality in whole-brain simulations emerges only within the critical regime (central panel) and only at the global coupling value corresponding to both global criticality and empirical fit (*G* = 0.028, Fig. 3-4).

Together, these findings demonstrate that the interplay between local criticality and global coupling shapes spatially structured gradients of timescales across the connectome, with *AC*1 modulation jointly determined by coupling strength and node in-strength, thereby enabling scale-free and flexible dynamics across scales.

## III Discussion

In this work, we introduce a multiscale, connectome-based modeling framework that unites the study of local (between neurons in a brain region) and global (between regions across the entire brain) criticality within a single formalism. To our knowledge, this provides the first mechanistic model displaying the coemergence of near-critical dynamics at both the local and global scales, and offers a new avenue for investigating multiscale mechanisms of brain dynamics. By tuning the brain regions to subcritical, critical, and supercritical regimes, and embedding them in the empirically derived mouse connectome, we explored how local regimes interact through long-range anatomical connections to shape both regional and whole-brain dynamics. Our results demonstrate that the relationship between local and global criticality is neither trivial nor one-directional: global coupling provides an extrinsic modulation that shifts regional dynamics toward or away from criticality depending on their initial state [33], and conversely, the tuning of local regimes (here fixed as homogeneous) sets the conditions for the emergence/suppression of scale-free activity across the entire brain.

A key finding is that global signatures of criticality emerge most robustly when local populations are tuned near the critical regime and coupled within an optimal range of global coupling. In this configuration, both empirical functional connectivity (FC) and dynamic functional connectivity (dFC) are best reproduced, and avalanche statistics display scale-free behavior at both the local and global levels. Based on the idea that criticality implies maximal sensitivity and susceptibility [31, 34], this result suggests that the brain may benefit from enhanced functional properties in both neuronal communication and interactions between distributed areas. Thus, the local critical regime has the advantage of naturally reconciling local and global signatures of criticality within a single parameter setting. At the same time, we observed that subcritical and supercritical neural masses can also yield good correspondence with empirical FC, despite disrupting scale-free statistics. These regimes display distinctive features—such as emergent oscillations in the supercritical case (Fig. 5)—that may be relevant for understanding diverse physiological and pathological states.

At the regional level (Fig. 5), the effect of coupling is strongly modulated by structural in-strength, producing topological gradients of autocorrelation timescales that become nonlinear in the critical regime. These gradients link to the broader hypothesis that regions higher in the cortical hierarchy exhibit longer timescales [35, 36] due to their proximity to criticality [30]. Anatomical constraints have been proposed as key determinants of such gradients [37–40]. Our results extend this view by showing that both local regime tuning and long-range coupling strength can shape the observed temporal hierarchy. In the brain in vivo, distance from criticality is typically estimated under conditions in which regions are already interconnected. Consequently, it remains unclear to what extent timescale gradients and fluctuations around criticality are shaped by intrinsic cytoarchitectural properties, neuromodulation, excitatory/inhibitory balance, versus long-range interactions [41].

These findings have important implications for understanding the fluctuations around criticality observed experimentally in local neuronal populations [42]. In particular, our results suggest that such fluctuations may not solely reflect local circuit properties or noise, but can also arise from topdown modulation via long-range interactions. This provides a mechanistic interpretation of recent electrophysiological observations that local criticality is not static but ‘fluid,’ varying across time and brain states [42]. In our framework, local populations near criticality can be transiently shifted toward sub- or supercritical dynamics by changes in input from connected regions (Fig. 2), which in turn are shaped by the evolving global state and the global coupling value. Alternatively, such fluctuations could be modeled via heterogeneous or time-varying local parameters. In general, different brain regions are characterized by distinct distances from criticality [30, 43], and these distances may themselves be dynamically modulated across time, tasks, and brain states.

This perspective raises the possibility that the lowdimensional embedding of brain dynamics observed in many empirical datasets [44–46]–where only a few principal components capture most of the variance–might reflect the imprint of large-scale network coordination on local dynamics. If fluctuations in local criticality are partly driven by global coupling, then the same long-range interactions that produce coherent low-dimensional trajectories could also regulate the temporal proximity of each region to its critical point.

Looking forward, our multiscale computational framework provides a platform for systematically investigating how structural connectivity, coupling strength, and heterogeneity in local tuning interact to produce the rich multiscale brain dynamics observed in experiments (see also [47–49]).

As it stands, the model presents limitations, and could be extended in several directions. For example: i) introducing heterogeneous local parameters to more closely match empirical variability and test how mixed regimes impact the dynamical signatures of global criticality [50, 51]; ii) incorporating time-varying parameters to simulate how task demands or neuromodulation may dynamically regulate the interaction between scales [52– iii) exploring pathological alterations in either local or global regimes to predict how disruptions at one scale propagate to the other, providing mechanistic insights into disorders characterized by altered criticality (e.g., epilepsy [56], schizophrenia [57], Parkinson’s disease [58], brain lesions [59]); iv) linking with low-dimensional manifold analyses to directly test whether the principal components of local criticality fluctuations align with large-scale network modes [44].

An important next step is to further integrate the study of criticality across local and global scales, which have traditionally been investigated in relative isolation. Advances in largescale electrophysiology now make it possible to record spiking activity from distributed neuronal populations across both inter- and intracortical circuits, offering an unprecedented opportunity to directly relate single-unit dynamics to systems-level critical fluctuations. Such multiscale datasets will allow rigorous empirical tests of whether—and how—local signatures of criticality propagate to, constrain, or are modulated by global network interactions. In the accompanying study [60], we take a first step in this direction by analyzing simultaneous recordings of large neuronal populations across distant brain areas, providing an experimental counterpart to the theoretical framework developed here.

In conclusion, by embedding minimal models of local dynamics within the mouse connectome, we have shown that local and global criticality are dynamically intertwined—their relationship being shaped by both intrinsic tuning and the structure of long-range interactions. Such an unifying approach not only advances the theoretical understanding of brain criticality and its multiscale dynamics but also offers testable hypotheses for experiments capable of bridging microcircuit and systems-level observations.

## IV Methods

### A Model Rationale

We developed a multiscale connectome-based model to investigate local and global brain criticality within a unified framework. Building on the observation that isolated neuronal populations, such as those in organotypic cultures, exhibit spontaneous scale-free neuronal avalanches [7], we hypothesize that neuronal populations can display critical-like behavior when driven solely by noisy inputs in the absence of long-range connections, and that coupling such populations via the empirical structural connectome allows us to study the emergence of global coordination and its feedback on local regimes. The model was implemented in three steps (Fig. 1A): (1) modeling the activity of a region of interest (ROI) as a network of neurons operating near criticality; (2) deriving a mean-field approximation that captures the critical-like dynamics while reducing computational complexity; and (3) simulating wholebrain activity by coupling ROIs through an empirical mouse connectome.

### B Neuron-Level Simulations

We modeled each ROI as a network of *K* interconnected neurons, represented as active rotators governed by the Shinomoto-Kuramoto model [24]:

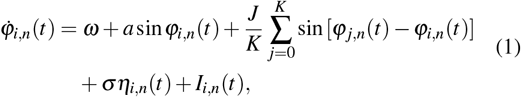

where *φ*_*i,n*_ is the phase of the *i*-th neuron in ROI *n* ∈ { 0, …, *M* }, *ω* = 1 is the intrinsic frequency, and *J* = 1.25 is the within-ROI all-to-all coupling strength. The population is driven by Gaussian white noise *η*_*i,n*_(*t*) of amplitude *σ*, and *a* is a *local* control parameter. Long-range coupling enters as an additive current:

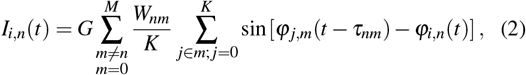

where *W*_*nm*_ is the structural connection weight between ROIs *n* and *m, τ*_*nm*_ is the propagation delay, and *G* is the *global coupling* parameter.

For *G* = 0, Eq. (1) describes a disconnected ROI. This model exhibits a hybrid-type (HT) synchronization transition [25, 26], where asynchronous states and incipient oscillations coexist, and where neuronal avalanche size and duration distributions span orders of magnitude [25]. For *G* ≠ 0, local dynamics are additionally shaped by long-range interactions according to the connectome.

### C Mean-Field Approximation

To reduce computational cost while retaining key dynamical properties, we adopted a mean-field description in terms of the Kuramoto order parameter:

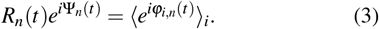

For the decoupled case (*G* = 0), Ott–Antonsen theory [61] yields the following mean-field equations [25, 26, 62]:

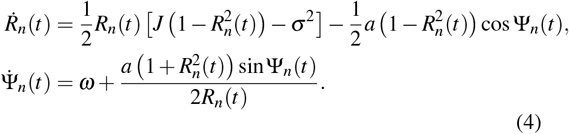

### D Identifying the Bistable Regime

We identified the bistable HT regime by scanning the (*a, σ*) parameter space of Eq. (4) (Suppl. Figure S1). For each parameter pair, we performed 250 deterministic simulations with different initial conditions, computed the time-averaged *R*_*n*_(*t*) for each run, and calculated the variance across initial conditions. This variance peaks in regions of bistability, where the final attractor depends on initial conditions. The resulting bistable region forms a triangular-shaped domain in (*a, σ*) space (Suppl. Figure S1).

### E Whole-Brain Mean-Field Simulations

The whole-brain model consists of 2*M* stochastic differential equations:

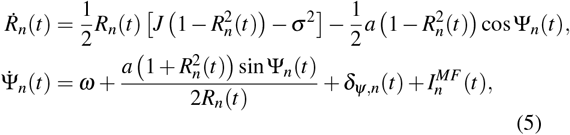

where we introduced a weak Gaussian noise term δ_*ψ,n*_(*t*) with standard deviation 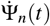 in Eq. (4) to emulate realistic fluctuations and recover avalanche-like dynamics in the bistable HT regime. The mean-field network input is:

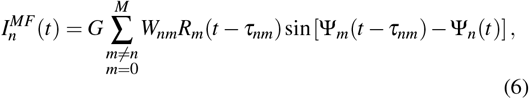

where *W*_*nm*_ are the weights of the structural connectome and we employed null delays *τ*_*nm*_ = 0, for simplicity.

### F Allen Mouse Structural Connectome

The structural connectivity matrix *W*_*nm*_ used in our simulations was derived from the tracer-based Allen Mouse Brain Connectivity Atlas [27], processed through the Virtual Mouse Brain pipeline [28], and previously employed in related work [59]. In these experiments, adult male C57Bl/6J mice received localized injections of a recombinant adeno-associated virus expressing an anterograde EGFP tracer. Tracer migration was imaged using serial two-photon tomography, revealing axonal projections from the injection site across the brain.

Injection density in each source region was defined as the fraction of infected pixels relative to the total number of pixels in that region. Projection density in a target region was quantified as the number of pixels carrying tracer signal, normalized by the total pixel count in that target. The directed connection strength from region *n* (source) to region *m* (target) was then computed as the ratio between projection density at *m* and injection density at *n*.

To construct the full connectome, we averaged the tracer data across injection experiments performed in the right hemisphere and considered projections to both ipsilateral and contralateral targets. This procedure yielded a weighted, directed connectivity matrix spanning 148 anatomical regions of interest (ROIs), including cortical and subcortical areas. For the present study, we restricted the analysis to the 60 cortical ROIs corresponding to our simulation parcellation (Suppl. Fig. S3). The final connectivity matrix was normalized such that all weights lie between 0 and 1.

### G Empirical Dataset

The data used in this study is part of a unified mouse resting-state functional MRI (rs-fMRI) resource collected by Dr. Joanes Grandjean and collaborators across two laboratories: ETH Zurich and the Singapore Bioimaging Consortium [63]. All data were obtained from the publicly available repository https://doi.org/10.34973/1he1-5c70 under the Creative Commons Attribution 4.0 International License (CC-BY 4.0). The datasets were preprocessed using a standardized pipeline to ensure comparability and consistency across experimental conditions.

To focus on a homogeneous control population, we included animals only if they were wild-type in genotype, classified as controls, scanned under the standardized medetomidine–isoflurane (mediso) protocol, and passed the quality assurance (QA) diagnostic tests (‘pass’). This yielded a final sample of 53 wild-type control animals (dataset label *CPS ctl* within the chronic social defeat, CSD1, collection).

Data acquisition was performed at a field strength of 9.4 Tesla using a cryogenically cooled phased-array receiver coil. rs-fMRI scans were acquired with TR = 1 s, TE = 9.2 ms, and consisted of 360 volumes per run, corresponding to a total acquisition time of 6 minutes. The mediso anesthesia protocol (medetomidine for sedation and low-dose isoflurane for maintenance) was used, as it minimizes physiological stress while preserving resting-state network activity.

The preprocessed time series were aligned to the Allen Institute for Brain Science (AIBS) reference atlas, resampled to a resolution of 0.2 × 0.2 × 0.2 mm^3^ (voxel volume 0.008 mm^3^). Out of the 604 regions defined in the AIBS atlas, we selected a subset of 60 cortical regions of interest (ROIs), corresponding to the parcellation used in our brain simulations (see Suppl. Fig. S2). Posterior cortical regions prone to imaging artifacts were excluded.

The resulting dataset therefore consisted of preprocessed rs-fMRI time series from 60 cortical ROIs across 53 animals, which served as the empirical benchmark for functional connectivity and dynamic analyses in comparison with simulated data (see next section).

### H Comparison with Empirical Data

Whole-brain model results correspond to simulated neuroelectric signals *R*_*n*_(*t*) at 2000 Hz time resolution for each brain region *n*, which were used for local and global criticality analyses. To compare our simulations to empirical data, these fast signals were convolved with a Balloon–Windkessel hemodynamic response function and z-scored, resulting in simulated BOLD (standardized) signals *B*_*n*_(*t*) at TR = 1s. Functional connectivity (FC) was computed as Pearson correlations between ROIs, and dynamic FC (dFC) from correlations between edge co-activation patterns *E*_*nm*_(*t*) = *B*_*n*_(*t*) *B*_*m*_(*t*) at different times *t*_*i*_ and *t* _*j*_ [64, 65]. For each *G*, we computed FC correlations and Kolmogorov–Smirnov distances between simulated and empirical FC/dFC distributions to identify the optimal *G*^⋆^ range.

## Acknowledgments

GR acknowledges support from the Marie Skłódowska–Curie Postdoctoral Fellowship (Project CAERUS) under the European Union’s Horizon Europe research and innovation programme (Grant No. 101199894). LDP acknowledges support from the European Union (ERC, NEMESIS, project number 101071900), AGAUR co-funded by the Departament de Recerca i Universitats de la Generalitat de Catalunya (AGAUR 2021 − SGR − 01165), and the Brazilian agency CNPq (Grant No. 444500*/*2024 − 3). Data were provided (in part) by the Donders Institute for Brain, Cognition and Behaviour, Radboud University Nijmegen.

## Author contributions

G.R. and L.D.P. designed research; G.R., P.B., D.D., and B.N. performed simulations; G.R. preprocessed experimental data; G.R., P.B., and L.D.P. analyzed data; G.R. and L.D.P. organized results; G.R. wrote the first draft; all authors discussed the results and wrote the paper.

## Code Availability

Full code for the reproduction of the simulations and data analysis described in this paper is freely available online at the repository https://github.com/grabuffo/Multiscale_Brain_Criticality.

## Supplementary Materials

**FIG. S1.**
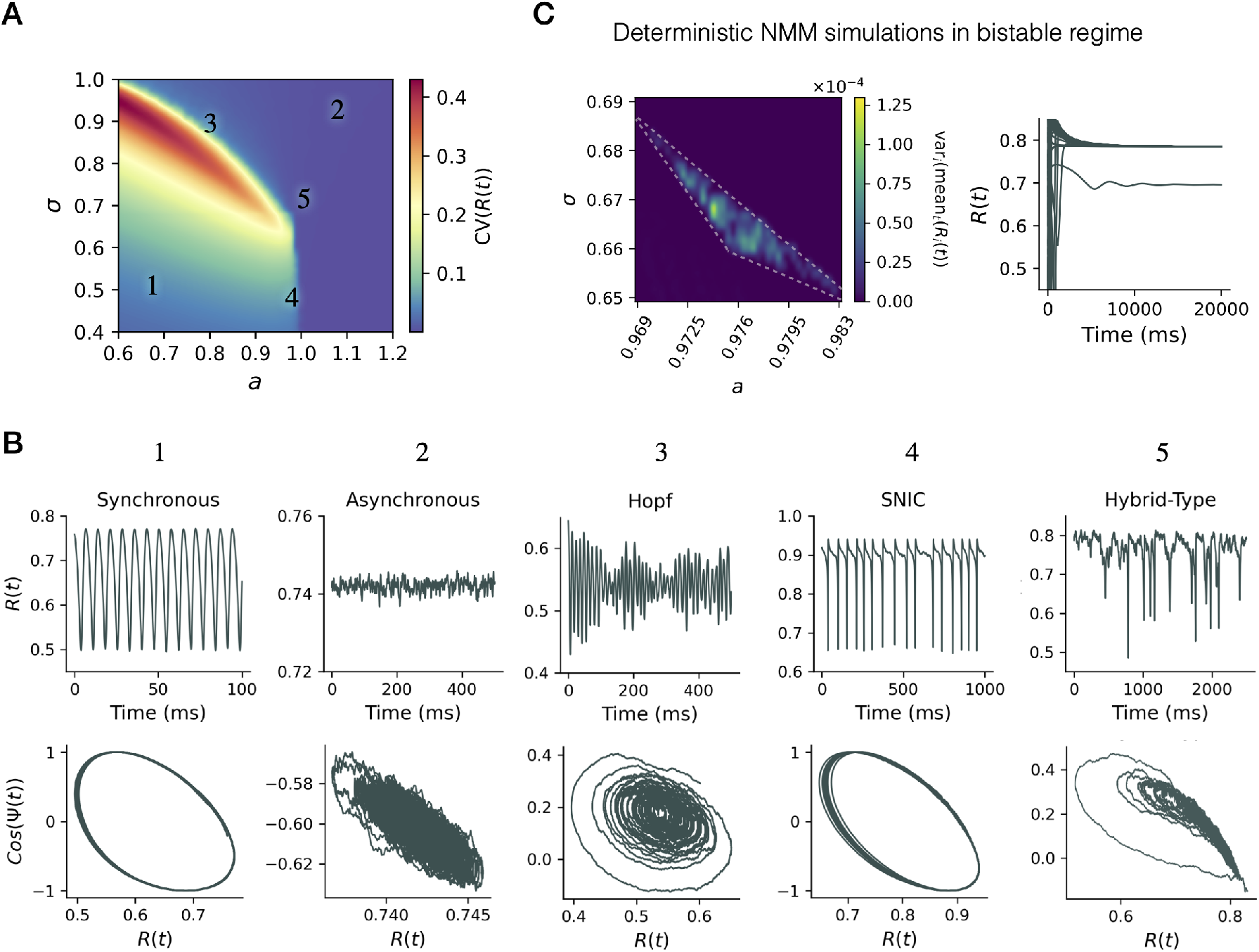
Local dynamical regimes of the mean-field neural mass model. **A)** Parameter space of the model as a function of *a* and *σ*, color-coded by the coefficient of variation of *R*(*t*). Five representative regimes are indicated: (1) synchronous oscillations, (2) asynchronous activity, (3) Hopf bifurcation, (4) saddle-node on invariant circle (SNIC), and (5) hybrid-type dynamics. For a corresponding schematic (non quantitative) representation of the mean-field model’s phase space, please refer to [25]. **B)** Examples of time series (top) and phase portraits (bottom, *R*(*t*) vs. cos Ψ(*t*) or *R*(*t*)) for the five regimes. **C)** Deterministic simulations in the bistable regime: left, variance-to-mean ratio of *R*(*t*) across *a* values highlights the coexistence of states; right, corresponding time series of *R*(*t*) showing convergence to different attractors depending on initial conditions.

**FIG. S2.**
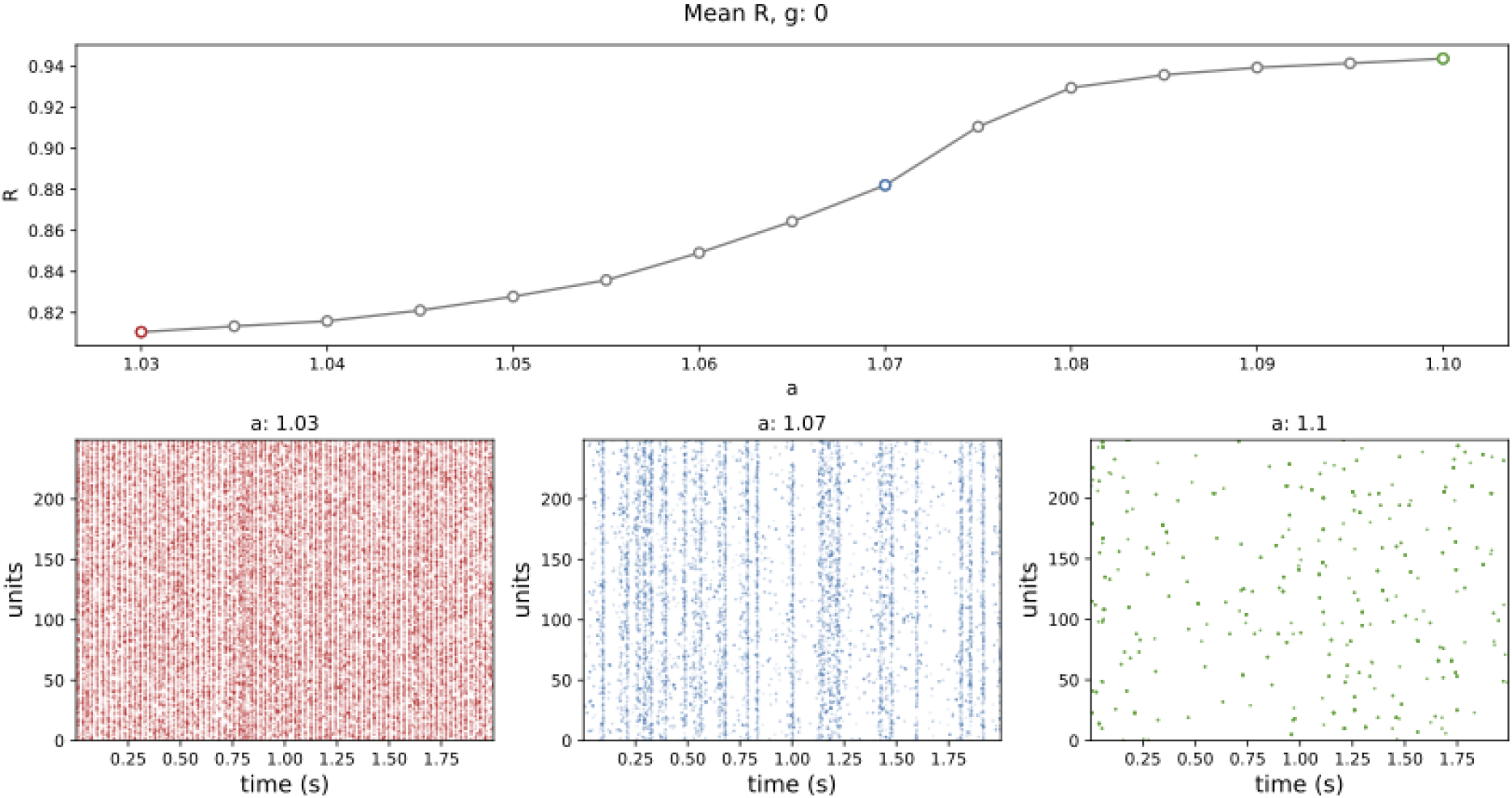
Single neuron simulations around the critical regime. (top) We simulated *N* = 5000 all-to-all connected active rotator neurons [24]. Following [25], we tuned the parameters around a hybrid-type bifurcation, fixing *σ* = 0.499, and tuning the parameter *a*, which allows us to obtain supercritical (red), critical (blue), and subcritical (green) regimes analogous to those obtained in Figure 1 using the meanfield model. As the parameter *a* increases, the system moves from low to high synchrony, as measured by the mean Kuramoto variable R. However, notice that the high synchrony of the subcritical regime corresponds to a low activity state. In other words, the neurons are coordinated in their inactivity. For more details on neuronal network simulations, see the accompanying paper [60]. Finally, notice that the correspondence between neuron-level and mean-field representations is not one-to-one in parameter space. In other words, we could not use the same parameters as [25] to locate the critical regime in the mean-field model. For that, we explored the (*a, σ*) parameter space in Figure 1

**FIG. S3.**
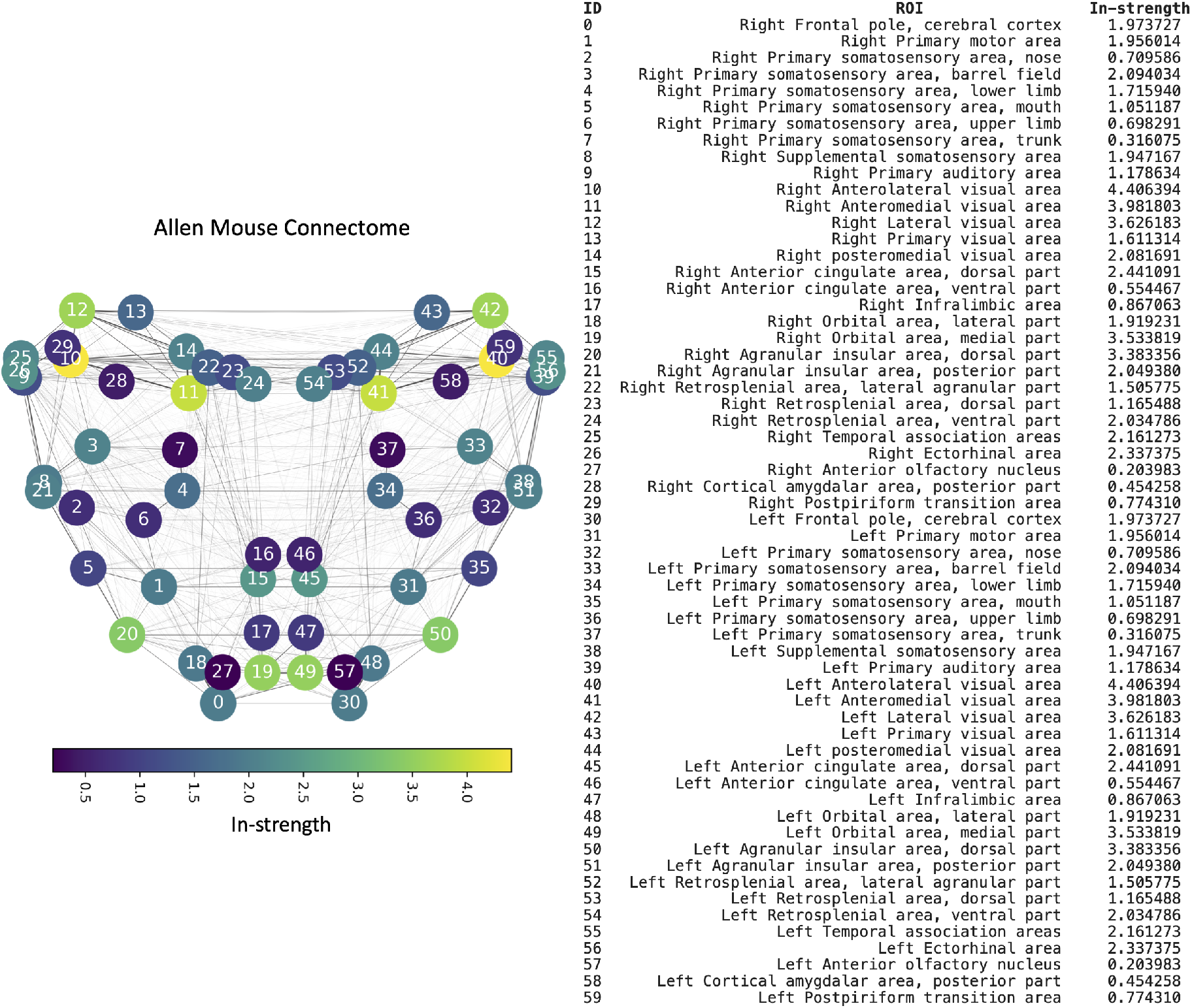
Allen Mouse Brain Connectome and regional in-strength. Left: Directed structural connectome of the Allen Mouse Brain Atlas represented as a circular layout of 60 cortical brain regions (30 per hemisphere). Node color encodes structural in-strength (sum of incoming weights), with warmer colors indicating higher in-strength values. Right: List of regions of interest (ROIs) with their corresponding ID and in-strength values.

## References

[1] John M Beggs. The criticality hypothesis: how local cortical networks might optimize information processing. Philosophical Transactions of the Royal Society A: Mathematical, Physical and Engineering Sciences, 366(1864):329–343, 2008.

[2] Luca Cocchi, Leonardo L Gollo, Andrew Zalesky, and Michael Breakspear. Criticality in the brain: A synthesis of neurobiology, models and cognition. Progress in neurobiology, 158: 132–152, 2017.

[3] Vincent Zimmern. Why brain criticality is clinically relevant: a scoping review. Frontiers in neural circuits, 14:565335, 2020.

[4] Jordan O’Byrne and Karim Jerbi. How critical is brain criticality? Trends in Neurosciences, 2022.

[5] Keith B Hengen and Woodrow L Shew. Is criticality a unified setpoint of brain function? Neuron, 113(16):2582–2598, 2025.

[6] Klaus Linkenkaer-Hansen, Vadim V Nikouline, J Matias Palva, and Risto J Ilmoniemi. Long-range temporal correlations and scaling behavior in human brain oscillations. Journal of Neuroscience, 21(4):1370–1377, 2001.

[7] John M Beggs and Dietmar Plenz. Neuronal avalanches in neocortical circuits. Journal of neuroscience, 23(35):11167–11177, 2003.

[8] W. L. Shew, H. Yang, S. Yu, R. Roy, and D. Plenz. Information capacity and transmission are maximized in balanced cortical networks with neuronal avalanches. J Neurosci, 31:55–63, Jan 2011. doi:10.1523/JNEUROSCI.4637-10.2011.

[9] W. L. Shew and D. Plenz. The functional benefits of criticality in the cortex. Neuroscientist, 19:88–100, Feb 2013. doi: 10.1177/1073858412445487.

[10] E. Tognoli and J. A. Kelso. The metastable brain. Neuron, 81: 35–48, 2014. doi:10.1016/j.neuron.2013.12.022.

[11] J. M. Beggs and D. Plenz. Neuronal avalanches in neocortical circuits. J Neurosci, 23:11167–11177, Dec 2003. doi: 10.1523/JNEUROSCI.23-35-11167.2003.

[12] J. M. Beggs and D. Plenz. Neuronal avalanches are diverse and precise activity patterns that are stable for many hours in cortical slice cultures. J Neurosci, 24:5216–5229, Jun 2004. doi:10.1523/JNEUROSCI.0540-04.2004.

[13] J. M. Palva, A. Zhigalov, J. Hirvonen, O. Korhonen, K. Linkenkaer-Hansen, and S. Palva. Neuronal long-range temporal correlations and avalanche dynamics are correlated with behavioral scaling laws. Proc Natl Acad Sci U S A, 110:3585–3590, Feb 2013. doi:10.1073/pnas.1216855110.

[14] G. M. Duma, A. Danieli, G. Mento, V. Vitale, R. S. Opipari, V. Jirsa, P. Bonanni, and P. Sorrentino. Altered spreading of neuronal avalanches in temporal lobe epilepsy relates to cognitive performance: A resting-state hdEEG study. Epilepsia, 64: 1278–1288, May 2023. doi:10.1111/epi.17551.

[15] O. Shriki, J. Alstott, F. Carver, T. Holroyd, R. N. Henson, M. L. Smith, R. Coppola, E. Bullmore, and D. Plenz. Neuronal avalanches in the resting MEG of the human brain. J Neurosci, 33:7079–7090, Apr 2013. doi:10.1523/JNEUROSCI.4286-12.2013.

[16] V. Priesemann, M. Valderrama, M. Wibral, and M. Le Van Quyen. Neuronal avalanches differ from wakefulness to deep sleep-evidence from intracranial depth recordings in humans. PLoS Comput Biol, 9(e1002985), Mar 2013. doi:10.1371/journal.pcbi.1002985.

[17] A. Zhigalov, G. Arnulfo, L. Nobili, S. Palva, and J. M. Palva. Relationship of fast- and slow-timescale neuronal dynamics in human MEG and SEEG. J Neurosci, 35:5385–5396, Apr 2015. doi:10.1523/JNEUROSCI.4880-14.2015.

[18] E. Tagliazucchi, P. Balenzuela, D. Fraiman, and D. R. Chialvo. Criticality in large-scale brain fMRI dynamics unveiled by a novel point process analysis. Front Physiol, 3, Feb 2012. doi: 10.3389/fphys.2012.00015.

[19] M. Yaghoubi, T. de Graaf, J. G. Orlandi, F. Girotto, M. A. Colicos, and J. Davidsen. Neuronal avalanche dynamics indicates different universality classes in neuronal cultures. Sci Rep, 8 (3417), Feb 2018. doi:10.1038/s41598-018-21730-1.

[20] Z. Bowen, D. E. Winkowski, S. Seshadri, D. Plenz, and P. O. Kanold. Neuronal Avalanches in Input and Associative Layers of Auditory Cortex. Front Syst Neurosci, 13, Sep 2019. doi: 10.3389/fnsys.2019.00045.

[21] George F Grosu, Alexander V Hopp, Vasile V Moca, Harald Bârzan, Andrei Ciuparu, Maria Ercsey-Ravasz, Mathias Winkel, Helmut Linde, and Raul C Mures, an. The fractal brain: scale-invariance in structure and dynamics. Cerebral Cortex, 33(8):4574–4605, 2023.

[22] Michelle Cirunay, Géza Ódor, István Papp, and Gustavo Deco. Scale-free behavior of weight distributions of connectomes. Physical Review Research, 7(1):013134, 2025.

[23] Markus Axer and Katrin Amunts. Scale matters: The nested human connectome. Science, 378(6619):500–504, 2022.

[24] Shigeru Shinomoto and Yoshiki Kuramoto. Phase transitions in active rotator systems. Progress of Theoretical Physics, 75(5): 1105–1110, 1986.

[25] Victor Buendía, Pablo Villegas, Raffaella Burioni, and Miguel A Munõz. Hybrid-type synchronization transitions: Where incipient oscillations, scale-free avalanches, and bistability live together. Physical Review Research, 3(2):023224, 2021.

[26] Vladimir Klinshov and Igor Franović. Two scenarios for the onset and suppression of collective oscillations in heterogeneous populations of active rotators. Physical Review E, 100 (6):062211, 2019.

[27] Seung Wook Oh, Julie A Harris, Lydia Ng, Brent Winslow, Nicholas Cain, Stefan Mihalas, Quanxin Wang, Chris Lau, Leonard Kuan, Alex M Henry, et al. A mesoscale connectome of the mouse brain. Nature, 508(7495):207–214, 2014.

[28] Francesca Melozzi, Marmaduke M Woodman, Viktor K Jirsa, and Christophe Bernard. The virtual mouse brain: a computational neuroinformatics platform to study whole mouse brain dynamics. Eneuro, 4(3), 2017.

[29] Marten Scheffer, Jordi Bascompte, William A Brock, Victor Brovkin, Stephen R Carpenter, Vasilis Dakos, Hermann Held, Egbert H Van Nes, Max Rietkerk, and George Sugihara. Earlywarning signals for critical transitions. Nature, 461(7260):53–59, 2009.

[30] Brendan Harris, Leonardo L Gollo, and Ben D Fulcher. Tracking the distance to criticality in systems with unknown noise. Physical Review X, 14(3):031021, 2024.

[31] O. Kinouchi and M. Copelli. Optimal dynamical range of excitable networks at criticality. Nature Phys, 2:348–351, 2006. doi:10.1038/nphys289.

[32] Adam Liska, Alberto Galbusera, Adam J Schwarz, and Alessandro Gozzi. Functional connectivity hubs of the mouse brain. Neuroimage, 115:281–291, 2015.

[33] Benedetta Mariani, Giorgio Nicoletti, Marta Bisio, Marta Maschietto, Stefano Vassanelli, and Samir Suweis. Disentangling the critical signatures of neural activity. Scientific reports, 12(1):10770, 2022.

[34] Woodrow L Shew and Dietmar Plenz. The functional benefits of criticality in the cortex. The neuroscientist, 19(1):88–100, 2013.

[35] John D Murray, Alberto Bernacchia, David J Freedman, Ranulfo Romo, Jonathan D Wallis, Xinying Cai, Camillo Padoa-Schioppa, Tatiana Pasternak, Hyojung Seo, Daeyeol Lee, et al. A hierarchy of intrinsic timescales across primate cortex. Nature neuroscience, 17(12):1661–1663, 2014.

[36] Adrián Ponce-Alvarez. Network mechanisms underlying the regional diversity of variance and time scales of the brain’s spontaneous activity fluctuations. Journal of Neuroscience, 45(10), 2025.

[37] Stefan J Kiebel, Jean Daunizeau, and Karl J Friston. A hierarchy of time-scales and the brain. PLoS computational biology, 4(11):e1000209, 2008.

[38] Rishidev Chaudhuri, Kenneth Knoblauch, Marie-Alice Gariel, Henry Kennedy, and Xiao-Jing Wang. A large-scale circuit mechanism for hierarchical dynamical processing in the primate cortex. Neuron, 88(2):419–431, 2015.

[39] Ryan V Raut, Abraham Z Snyder, and Marcus E Raichle. Hierarchical dynamics as a macroscopic organizing principle of the human brain. Proceedings of the National Academy of Sciences, 117(34):20890–20897, 2020.

[40] Junhao Liang, Zhuda Yang, and Changsong Zhou. Excitation– inhibition balance, neural criticality, and activities in neuronal circuits. The Neuroscientist, 31(1):31–46, 2025.

[41] Eli J Müller, Brandon R Munn, and James M Shine. The brain that controls itself. Current Opinion in Behavioral Sciences, 63:101499, 2025.

[42] Antonio J Fontenele, Nivaldo AP De Vasconcelos, Thaís Feliciano, Leandro AA Aguiar, Carina Soares-Cunha, Bárbara Coimbra, Leonardo Dalla Porta, Sidarta Ribeiro, Ana João Rodrigues, Nuno Sousa, et al. Criticality between cortical states. Physical review letters, 122(20):208101, 2019.

[43] Guillermo B Morales, Serena Di Santo, and Miguel A Munõz. Quasiuniversal scaling in mouse-brain neuronal activity stems from edge-of-instability critical dynamics. Proceedings of the National Academy of Sciences, 120(9):e2208998120, 2023.

[44] Antonio J Fontenele, J Samuel Sooter, V Kindler Norman, Shree Hari Gautam, and Woodrow L Shew. Low-dimensional criticality embedded in high-dimensional awake brain dynamics. Science Advances, 10(17):eadj9303, 2024.

[45] Annie E Cathignol, Lionel Kusch, Marianna Angiolelli, Emahnuel Troisi Lopez, Arianna Polverino, Antonella Romano, Giuseppe Sorrentino, Viktor Jirsa, Giovanni Rabuffo, and Pierpaolo Sorrentino. Magnetoencephalography dimensionality reduction informed by dynamic brain states. European Journal of Neuroscience, 61(9):e70128, 2025.

[46] Matthew G Perich, Devika Narain, and Juan A Gallego. A neural manifold view of the brain. Nature Neuroscience, 28(8): 1582–1597, 2025.

[47] Sheng-Jun Wang, Claus C Hilgetag, and Changsong Zhou. Sustained activity in hierarchical modular neural networks: selforganized criticality and oscillations. Frontiers in computational neuroscience, 5:30, 2011.

[48] Roberta Russo, Hans Jürg Herrmann, and Lucilla De Arcangelis. Brain modularity controls the critical behavior of spontaneous activity. Scientific Reports, 4(1):4312, 2014.

[49] Samora Okujeni and Ulrich Egert. Structural modularity tunes mesoscale criticality in biological neuronal networks. Journal of Neuroscience, 43(14):2515–2526, 2023.

[50] Gustavo Deco, Morten L Kringelbach, Aurina Arnatkeviciute, Stuart Oldham, Kristina Sabaroedin, Nigel C Rogasch, Kevin M Aquino, and Alex Fornito. Dynamical consequences of regional heterogeneity in the brain’s transcriptional landscape. Science Advances, 7(29):eabf4752, 2021.

[51] Leonardo Dalla Porta, Jan Fousek, Alain Destexhe, and Maria V. Sanchez-Vives. Cholinergic heterogeneity facilitates synchronization and information flow in a whole-brain model. bioRxiv, 2025. doi:10.1101/2025.10.28.685048. URL https://www.biorxiv.org/content/early/2025/10/29/2025.10.28.685048.

[52] Mojtaba Nazari, Javad Karimi Abadchi, Milad Naghizadeh, Edgar J Bermudez-Contreras, Bruce L McNaughton, Masami Tatsuno, and Majid H Mohajerani. Regional variation in cholinergic terminal activity determines the non-uniform occurrence of cortical slow waves during rem sleep in mice. Cell reports, 42(5), 2023.

[53] Brandon R Munn, Eli J Müller, Gabriel Wainstein, and James M Shine. The ascending arousal system shapes neural dynamics to mediate awareness of cognitive states. Nature communications, 12(1):6016, 2021.

[54] Richard Gao, Ruud L Van den Brink, Thomas Pfeffer, and Bradley Voytek. Neuronal timescales are functionally dynamic and shaped by cortical microarchitecture. elife, 9:e61277, 2020.

[55] Roxana Zeraati, Yan-Liang Shi, Nicholas A Steinmetz, Marc A Gieselmann, Alexander Thiele, Tirin Moore, Anna Levina, and Tatiana A Engel. Intrinsic timescales in the visual cortex change with selective attention and reflect spatial connectivity. Nature communications, 14(1):1858, 2023.

[56] Shun Liu, Fali Li, and Feng Wan. Distance to criticality undergoes critical transition before epileptic seizure attacks. Brain Research Bulletin, 200:110684, 2023.

[57] Golnoush Alamian, Tarek Lajnef, Annalisa Pascarella, Jean-Marc Lina, Laura Knight, James Walters, Krish D Singh, and Karim Jerbi. Altered brain criticality in schizophrenia: new insights from magnetoencephalography. Frontiers in Neural Circuits, 16:630621, 2022.

[58] Pierpaolo Sorrentino, Rosaria Rucco, Fabio Baselice, Rosa De Micco, Alessandro Tessitore, Arjan Hillebrand, Laura Mandolesi, Michael Breakspear, Leonardo L Gollo, and Giuseppe Sorrentino. Flexible brain dynamics underpins complex behaviours as observed in parkinson’s disease. Scientific reports, 11(1):4051, 2021.

[59] Giovanni Rabuffo, Houefa-Armelle Lokossou, Zengmin Li, Abolfazl Ziaee-Mehr, Meysam Hashemi, Pascale P Quilichini, Antoine Ghestem, Ouafae Arab, Monique Esclapez, Parul Verma, et al. Mapping global brain reconfigurations following local targeted manipulations. Proceedings of the National Academy of Sciences, 122(16):e2405706122, 2025.

[60] Leonardo Dalla Porta, Pietro Bozzo, Marco N. Pompili, Damien Depannemaecker, Antonio J. Fontenele, Tomoki Fukai, Pierpaolo Sorrentino, and Giovanni Rabuffo. Intra- and interhemispheric signatures of criticality at the onset of synchronization. BioRxiv, 2025.

[61] Edward Ott and Thomas M Antonsen. Low dimensional behavior of large systems of globally coupled oscillators. Chaos: An Interdisciplinary Journal of Nonlinear Science, 18(3), 2008.

[62] VV Klinshov, D. Zlobin, BS Maryshev, and DS Goldobin. Effect of noise on the collective dynamics of a heterogeneous population of active rotators. Chaos: An Interdisciplinary Journal of Nonlinear Science, 31(4), 2021.

[63] Joanes Grandjean. A common mouse fmri resource through unified preprocessing. 2020.

[64] Giovanni Rabuffo, Jan Fousek, Christophe Bernard, and Viktor Jirsa. Neuronal cascades shape whole-brain functional dynamics at rest. eneuro, 8(5), 2021.

[65] Maria Pope, Makoto Fukushima, Richard F Betzel, and Olaf Sporns. Modular origins of high-amplitude cofluctuations in fine-scale functional connectivity dynamics. Proceedings of the National Academy of Sciences, 118(46):e2109380118, 2021.

